# Methods to Reduce Sea Turtle Interactions in the Atlantic Canadian Pelagic Long Line Fleet

**DOI:** 10.1101/117556

**Authors:** Zachary T. Sherker

**Affiliations:** St. Francis Xavier University, Department of Biology, Antigonish, Nova Scotia B2G 2W5, Canada

## Abstract

This project investigates the role of fisheries management in the conservation of loggerhead (*Caretta caretta*) and leatherback sea turtles (*Dermochelys coriacea),* both of which are currently listed as vulnerable by the IUCN (International Union for Conservation of Nature). These species migrate from nesting grounds in South America to feed on gelatinous zooplankton (jellyfish) in the North Atlantic off the coast of the United States and Canada. The seasonal foraging grounds of sea turtles heavily overlap with areas of high fishing effort for the longline tuna and swordfish fleet, a fishery that has significantly high rates of sea turtle incidents. The dynamic nature of sea turtle foraging patterns renders static spatio-temporal fishing area closures ineffective. Rather, turtle by-catch mitigation requires small-scale, event-triggered closures and decentralized management to reduce incidents while minimizing the negative socio-economic impact of area closures on fishermen. A number of methods that increase fishing selectivity have been implemented in other commercial fisheries around the globe and are suggested for the Atlantic Canadian fleet moving forward.

## 1. Introduction

The two most common species of sea turtle found in Atlantic Canadian waters are the leatherback sea turtle, *Dermochelys coriacea,* and the loggerhead sea turtle, *Caretta caretta* (McAlpine *et al.* 2007). Although there is seasonal variability in the number of turtles located off the coast of eastern Canada, recent studies suggest that northern foraging grounds are critical habitat for western North Atlantic subpopulations (McAlpine *et al.* 2007: Fossett *et al.* 2010: James *et al.* 2005).

### 1.1 Leatherback Presence in Canadian Waters

Leatherback sea turtles undergo an annual migration from nesting beaches in the southern United States and South America to feed exclusively on gelatinous zooplankton (i.e. scyphozoan jellyfish, *Cyanae capillata*) at higher latitudes (James & Herman 2001: Fossette *et al.* 2010). Leatherbacks are present in Canadian waters from June to November (James *et al.* 2005b) with abundance peaking between August and September (McAlpine *et al.* 2007). Arrival in northern foraging grounds appears to coincide with seasonal oceanic conditions that favor gelatinous zooplankton blooms (Sherrill-Mix *et al.* 2008:Witt *et al.* 2007). Due to the low nutritional value of their prey, leatherbacks need to consume an average of 60-120kg of gelatinous zooplankton per day (Jones *et al.* 20121 and are continuously foraging during daylight hours (Wallace *et al.* 2015).

The frequency of leatherback sightings tends to increase with rising sea surface temperatures (McAlpine *et al.* 2007). This increase is likely a result of the favorable energetic efficiency of foraging in warmer conditions, as opposed to a thermal restriction on the turtles (James & Mrosovsky 2004:Wallace *et al.* 2015:Casey *et al.* 2014). Physiological adaptations and the large body size of leatherback sea turtles allows individuals to forage in cooler waters that often induce lethargy in other marine turtle species (Witt *et al.* 2007:Greer 1973: James *et al.* 2006). although dives below the thermocline in Canadian waters are rare due to the shallow distribution of the prey species (Wallace *et al.* 2015:Fossette *et al.* 2010:Hays 2008: Fernández-Álamo & Färber-Lorda 2006: Bailev *et al.* 2012:Hamelin *et al.* 2014).

All leatherback sea turtles observed in Canada have been classified as adults or large sub-adults (James *et al.* 2005b:McAlpine *et al.* 2007) and their size generally increases as we move north into cooler waters (Witt *et al.* 2007:James *et al.* 2007; Dodge *et al.* 2014; James *et al.* 2005). Lower lipid stores and a relatively small body size may restrict most sub-adults and juveniles from migrating as far north as Canada (Dodge *et al.* 2014). If climate change results in increased sea surface temperatures, the foraging habitat of leatherbacks may expand northward. This would result in more individuals as well as smaller individuals foraging in Canadian waters in years to come (McMahon & Hays 2006).

### 1.2 Loggerhead Presence in Canadian Waters

Loggerhead sea turtles exhibit a biphasic life history consisting of an oceanic (offshore) juvenile stage and a neritic (inshore) adult stage (Bolton 2003). As hatchlings from northwest Atlantic nesting beaches transition to juveniles, they are picked up by the Gulf Stream and are passively carried to higher latitudes, where they enter the oceanic zone (>200m in depth) (Bolton 2003; Harris *et al.* 2010). Juvenile loggerheads forage year-round in the oceanic zone (Hochscheid *et al.* 2007). moving north into Canadian waters during the spring and summer months, when productivity is high and sea surface temperatures begin to rise, and retreating closer to the Gulf Stream for fall and winter (Mansfield *et al.* 2009).

The diet of loggerhead sea turtles in Canadian waters is largely unknown (Harris *et al.* 2010). although feeding behaviors can be inferred from studies of the mid-North Atlantic, eastern North Atlantic, and North Pacific populations. Juvenile loggerheads are epipelagic (top 200m of the oceanic zone) foragers that primarily exhibit short, shallow dives of 2-5 meters, and spend approximately 75% of their time in the top 5m of the water column (Bolton 2003). Oceanic loggerheads in the North Atlantic feed on a variety of epipelagic prey, especially *Cyanea capillata* and salps (planktonic tunicates) (Smolowitz *et al.* 2015: Harris *et al.* 2010).

Juvenile loggerheads forage for 10-18 years in the oceanic zone (Harris *et al.* 2010). moving both passively and actively relative to surrounding water currents (Bolton 2003: Putman & Mansfield 2015). The role that the Gulf Stream plays in the distribution of juvenile loggerheads can explain their commonality in Canadian waters east of the continental shelf (McAlpine *et al.* 2007). Loggerheads are poor thermoregulators and can only maintain a body temperature 1-2°C above surrounding water temperatures (Zug *et al.* 2001). As such, their distribution is largely limited by water temperature (Braun-McNeill *et al.* 2008). There is a significant correlation between size distribution and latitude in juvenile loggerheads, with larger individuals observed at higher latitudes (Arendt *et al.* 2012). suggesting that body mass is the primary physiological component that drives thermoregulation for this species. Juvenile loggerheads are strongly associated with sea surface temperatures (SST) above 15°C (Narazaki *et al.* 2015: Hochscheid *et al.* 2007) and have never been caught by the Canadian pelagic long-line fleet below this thermal threshold (Brazner & McMillan 2008). Between 5.0-9.0°C, juveniles undergo cold stunning, which can result in lethargy, a loss of buoyancy, inability to dive, and reduced feeding (Shwartz 1978). although individuals have been observed actively foraging in temperatures as low as 7°C (Smolowitz *et al.* 2015). Due to the cool summer SST of coastal Canadian waters (12-13°C) (McAlpine *et al.* 2007). Loggerheads are only occasionally observed inshore (Brazner & McMillan 2008). An exception to this is when warm-core rings break off of the Gulf Stream and carry warm water from the Sargasso Sea northwest, creating temporary ‘hotspots’ in coastal waters (McAlpine *et al.* 2007). Since these oceanic processes are relatively unpredictable, the presence of juvenile loggerheads close to shore is considered highly variable (McAlpine *et al.* 2007). If sea surface temperatures rise, the forging habitat of loggerhead sea turtles could expand further north into Canadian waters (Witt *et al.* 2010).

### 1.3 Current Status

The global populations of both leatherback sea turtles and loggerhead sea turtles are currently listed as “vulnerable” in the International Union on the Conservation of Nature (IUCN) Red List (Wallace *et al.* 2013; Casale & Tucker 2015). Leatherbacks are considered “endangered” by Canada’s Species at Risk Act (SARA). The United States’ Endangered Species Act (ESA) considers leatherbacks “endangered” throughout its range and lists the northwest Atlantic subpopulation of loggerheads as “threatened” (NOAA 2014). The Committee on the Status of Endangered Wildlife in Canada (COSEWIC) designated the Atlantic leatherback as “endangered” under Criterion A due to global population decline in 2012 (COSEWIC 2012).

A 28% decline in the loggerhead nesting populations of the Peninsular Florida Recovery Unit (accounts for 80% of all loggerhead nesting in the Atlantic) was recorded between 1989 and 2006 (Witherington *et al.* 2009). The steep negative trend of northwest Atlantic nesting populations provided rationale for the designation of loggerheads as “endangered” under Criterion A of COSEWIC in 2010 (COSEWIC 2010).

It should be noted that trends in loggerhead nesting populations are not a reliable indicator of the recent impact of fisheries by-catch on the population since a significant decline in juveniles may not translate to a decline in nesting populations for 10-20 years or more (Ehrhart *et al.* 2014). Also, extensive mixing of genetically distinct loggerhead populations occurs in oceanic waters (Bowan *et al.* 2005; Bowan & Karl 2007) and as a result a high level of overall juvenile loggerhead mortality during the oceanic stage may result in only negligible changes in the nesting trends of various sub-populations.

### 1.4 International Protection

Loggerheads and Leatherbacks are protected under Appendix I of the Convention on International Trade in Endangered Species of Wild Fauna and Flora (CITES), which renders commercial harvest illegal. Canada has been a signatory of CITES since 1975. Both loggerhead and leatherback sea turtles are protected under Appendix I and II of the Convention on the Conservation of Migratory Species of Wild Animals (CMS). No North American country is a member state of CMS. The Inter-American Convention for the Protection and Conservation of Sea Turtles (IAC) is an international instrument that promotes the protection of sea turtle populations and associated high-use habitats (IAC 2004). Canada is one of only four western countries with an Atlantic coastline that is not a member of IAC.

> “Canada is the only western country with a marine coastline that does not have any federal legislation directed at marine sea turtles specifically” (McApline *et al.* 2007).

### 1.5 Influence of Fisheries By-catch

Fisheries by-catch is the primary cause for population decline in all species of marine turtle and is considered the biggest threat to the population recovery of both loggerheads and leatherbacks (Lewison *et al.* 2004:Wallace *et al.* 2010:Wallace *et al.* 2011: Wallace *et al.* 2013). Long line, gillnet, and trawl fishing gear have the highest turtle by-catch rates observed in commercial fisheries, with gillnet and trawl having significantly higher mortality rates than long line (Wallace *et al.* 2013; Lewison *et al*. 2013). although this analysis does not consider the potentially high post-release mortality associated with long line-turtle interactions (Garrison & Stokes 2014; Chaloupka *et al.* 2004; Parker *et al.* 2005; Sasso & Epperly 2007; Hays *et al.* 2003; Ryder *et al.* 2006; Swimmer *et al.* 2006; Swimmer *et al.* 2014). Turtle Excluder Devices (TED) with large escape openings have been implemented in trawl fisheries around the world and have drastically reduced the by-catch and associated mortality of turtles by trawling without significantly impacting target catch rates (Epperly 2003; Brazner & McMillan 2008; Finkbeiner *et al.* 2011). The recent Marine Stewardship Council (MSC) certification received by the Canadian herring gillnet fishery indicates that gillnets are not a significant source of mortality for sea turtles in Canadian waters.

Turtles by-catch by the pelagic long line fleet directed at tuna and swordfish is the only direct anthropogenic source of mortality for loggerheads and leatherbacks currently recognized in Canadian waters (Harris *et al.* 2010; DFO 2010). although entanglement in fixed gear (e.g. lobster and crab pots) in coastal waters may be a significant source of mortality that is overlooked due to the relative rarity of interaction events (James *et al.* 2005a). A conservative estimate quantifying the number of marine turtles caught globally by long line gear suggests that at least 220,000 loggerheads and 50,000 leatherbacks were either hooked or entangled in gear in the 2000 fishing season alone, with 37% of all long line fishing effort occurring in the Atlantic Ocean (Lewison *et al.* 2004). Canadian waters are considered a high threat area for both loggerheads and leatherbacks as a result of the high spatio-temporal overlap of the Canadian Pelagic Long Line Fishery with critical foraging habitat for marine turtles (Wallace *et al.* 2011:Wallace *et al.* 2013).

The Canadian Pelagic Long Line Fleet directed for swordfish and “other tuna” (i.e yellowfin (*Thunnus albacares*), bigeye (*Thunnus obesus*), and albacore (*Thunnus alalunga*)) caught approximately 1,200 loggerheads annually between 2002-2008 (Brazner & McMillan 2008). with higher by-catch rates observed when the gear was targeting tuna (Paul *et al.* 2010; Brazner & McMillan 2008). A similar magnitude of marine turtle by-catch is exhibited in the U.S. and Canadian Pelagic Long Line Fisheries, likely due to the close proximity of their fishing efforts (Brazner & McMillan 2008). The U.S. Atlantic Pelagic Long Line Fleet caught approximately 365 leatherback turtles in 2013 (Garrison & Stokes 2014). It can be inferred that a similar number of incidents occurred in Atlantic Canadian waters during the same fishing season, although mortality of leatherback sea turtles in Canadian waters is considered to be low (Atlantic Leatherback Turtle Recovery Team 2006).

There may be a higher post-release survival rate for leatherbacks compared to loggerheads due to the nature of hooking incidents. Leatherbacks are primarily found either entangled in long line gear or externally hooked, whereas loggerheads are almost exclusively hooked internally (Garrison & Stokes 2014). The relatively large body size of Leatherbacks may account for their increased susceptibility to entanglement (Atlantic Leatherback Turtle Recovery Team 2006).

## 2. Potential Sea Turtle By-catch Mitigation Measures for Long Line Fisheries

The primary goals of a successful mitigation method is to minimize any ecosystem degradation caused by fishing operations while maintaining the economic viability of the fishery being managed (Dunn *et al.* 2011; O’Keefe *et al.* 2013). The most efficient way to achieve both objectives is through increasing the fishing selectivity (i.e. amount of target catch per by-catch event) of active vessels (Dunn *et al.* 2011). The most common approaches to reducing by-catch include the static area closure of critical habitat, annual by-catch caps, fleet communication to identify and avoid areas with high by-catch rates, and gear modifications (O’Keefe *et al.* 2013). Measures that minimize by-catch by means of fishing effort reductions (e.g. long-term closures and by-catch caps) do not impact fishing selectivity (Dunn *et al.* 2011) and in turn levy unwanted economic impacts on fishermen (O’Keefe *et al.* 2013.). These measures should therefore be considered a last ditch attempt to alleviate negative fisheries impact after all other measures fail. On the contrary, modifications to operational procedures (e.g. fleet communication, dynamic closures, and gear restrictions) have the potential to increase fishing selectivity and lessen negative economic impacts of by-catch mitigation on fishermen (Dunn *et al.* 2011).

### 2.1 Marine Protected Areas/ Long-term fishing closures

Marine protected areas (MPA) are areas of the ocean in which human activities (e.g. fishing) are restricted in order to conserve critical marine habitats and maintain biodiversity (Edgar *et al.* 2007). MPAs and long-term fishing closures levy a similar constraint to fishing operations and will therefore be analyzed as a single management technique for the purposes of this paper. The Canadian federal government has recently committed to raising the proportion of Canadian coastal and oceanic waters that are considered protected areas from the current 1.3% to 5% by 2017 and 10% by 2020, in accordance with targets set at the International Convention on Biodiversity in 2010 (Liberal Party of Canada 2016). The oceanic conditions that govern sea turtle movements and aggregations are highly dynamic (Roe *et al.* 2014; Howell *et al.* 2008; Witt *et al.* 2007; Dunn *et al.* 2011). whereas the space allocated by MPAs and long-term area closures is static. The mismatch of the basic nature of long-term closures and critical sea turtle habitat in Canadian waters renders MPAs an ineffective measure for reducing sea turtle by-catch in the long line fleet (Game *et al.* 2009).

Long-term area closures directed at the protection of sea turtles can displace fishing effort into areas that provide habitat for other species of concern (Baum *et al.* 2003). therefore this mitigation method fails to reduce the impact of by-catch on biodiversity but rather redistributes it amongst other susceptible species (O’Keefe *et al.* 2013). Furthermore, much of this displaced fishing effort is highly localized around MPA borders (Murawski *et al.* 2005). potentially forming a wall of fishing effort that sea turtles would have to travel through in order to access the critical habitat being protected. Long-term closures can also have negative economic impacts on fishermen, including longer steam times to arrive at fishing grounds that are further from shore and reduced catch rates of the target species (O’Keefe *et al.* 2013).

### 2.2 By-catch Caps

Annual by-catch caps are limits placed on the number of incidents a fishery can have with a specific non-target species within a fishing season. Once the by-catch limit is reached, the fishing season closes, resulting in reduced target species landings and substantial economic loss for fishermen. Annual incidental catch quotas for loggerheads and leatherbacks introduced in some long line fisheries in the U.S. (FAO 2009) have had little impact on slowing population decline. It is likely that without 100% observer coverage to monitor by-catch and enforce limits, incentives for fishermen to not report incidents and prolong the fishing season will be high (O’Keefe *et al.* 2013). Fishermen will also be less likely to follow proper bycatch handling protocol if sea turtles are identified as the limiting factor to their economic pursuits, potentially reducing post-incident survival of turtles caught in long line gear.

### 2.3 Voluntary Closures/ By-catch “Hotspot” Avoidance

Voluntary closures that are either event-or temperature-triggered can reduce sea turtle interactions by increasing fishing selectivity and directing fishing efforts into areas with lower levels of by-catch per unit of effort (BPUE) (Dunn *et al.* 2011; Gilman *et al.* 2006; Howell *et al.* 2008; Howell *et al.* 2015). Event-triggered closures exploit the fact that there is a higher probability of a sea turtle interaction occurring in the days following an initial by-catch event in that area (Gardner *et al.* 2008; Gilman *et al.* 2007; Lewison *et al.* 2009) since sea turtles are often foraging when they are caught on long line gear (Watson *et al.* 20051 and aggregations of marine predators in the oceanic zone are largely driven by prey availability (Davoren 2013; Houghton *et al.* 2006; Wallace *et al.* 2015:Bailey *et al.* 2012; Dewar *et al.* 2011). Therefore, if a sea turtle is caught on gear, it is likely that the gear is set in an area with high prey availability. Real-time vessel communication to identify areas with high rates of sea turtle captures, via email or radio, is suspected to reduce turtle bycatch in the U.S. Atlantic swordfish fleet by about 50% (Gilman *et al.* 2006).

On the other hand, voluntary closures based on sea surface temperature allow fishermen to avoid sea turtles prior to placing any hooks in the water. Sea surface temperature is used as a gage to initiate rolling closures in some U.S. Atlantic fisheries in order to avoid interactions with sea turtles (Murray 2009). The TurtleWatch program in the Hawaiian pelagic long line fleet provides fishermen with up-to-date maps that identify areas where vessels are likely to encounter loggerhead and leatherback sea turtles, based on the current thermal conditions (Howell *et al.* 2008; Howell *et al.* 2015). In the first fishing season following the introduction of the TurtleWatch program, 67% of all loggerhead by-catch events occurred in areas that were identified by the program as having a high probability of sea turtle presence (Howell *et al.* 2008). Through collaboration between scientists and industry, the TurtleWatch product provides fishermen with the tools to evade sea turtles while maintaining a high target catch rate (Howell *et al.* 2015). Both fleet communication and by-catch hotspot avoidance are relatively inexpensive mitigation methods that may help to separate fishing effort from sea turtle habitat in time without the need for direct regulation from resource managers (Gilman *et al.* 2006:Howell *et al.* 2008; Howell *et al.* 2015).

### 2.4 Modifying Fishing Practices

The modification to long line fishing practices that has been most thoroughly investigated for reducing sea turtle interactions is lowering the depth at which baited hooks are set. This is often done by elongating the float line (line that connects floats to the mainline) and using a weight to drag all hooks below the desired depth (FAO 2009; Beverly *et al.* 2009). Modifying fishing practices to shift effort into waters deeper than 40 meters would likely result in a reduction of sea turtle by-catch in Canadian waters, given that sea turtles rarely dive beneath the thermocline in these regions (Bailey *et al.* 2012; Wallace *et al.* 2015; Bolton 2003). Furthermore, there is little historical by-catch by the Canadian pelagic long line fleet below 40 meters (Brazner & McMillan 2008). which aligns with the results of experimental data that suggests that the highest rate of turtle by-catch occurs within the top 50 meters of the water column (Gilman *et al.* 2006; Beverly *et al.* 2009).

Lowering the depth of long line fishing operations would not be feasible for target species that are fished at shallow depths (e.g. swordfish (FAO 2009)) (Brazner & McMillan 2008). but would be expected to have little impact on the catch of deeper species (e.g. tuna (Dagorn *et al.* 2000)) (Beverly *et al.* 2009; Polovina *et al.* 2003). The only additional cost associated with fishing in deeper waters is the increased time required for gear deployment and retrieval (Beverly *et al.* 2009). although this cost would be offset by a reduction in the time spent removing hooks and line from turtles if by-catch were successfully reduced. With fewer turtles caught in the gear, there will also be more hooks free to catch the target species (Cox *et al.* 2007).

### 2.5 Modifying Fishing Gear/Bait

The most common methods aimed at reducing sea turtle by-catch and mortality in long line fisheries include modifications to the hook type, bait species, and the technique used to hook the bait. Switching from the traditional J-hook with squid bait to the wider circle-hook with mackerel bait has shown to significantly reduce loggerhead by-catch by as much as 90% (Gilman *et al.* 2007; Watson *et al.* 20051. while increasing the catch of swordfish by 30% (Watson *et al.* 2005). Leatherback by-catch was most successfully abated by using a J-hook and mackerel bait, with swordfish catch under this treatment increasing by 67% compared to J-hooks baited with squid (Watson *et al.* 2005). Swordfish and tuna catch rates display highly dissimilar responses to various hook-bait combinations (Watson *et al.* 2005). suggesting that these modifications should be further examined within species-specific fisheries to avoid undesirable effects on target species catch rates.

Although the use of circle-hooks have been a promising method to reduce sea turtle by-catch in some long line fisheries (Santos *et al.* 2013; Watson *et al.* 2005; Sales *et al.* 2010; Gilman *et al.* 2007). there is variability in its effectiveness in Atlantic fleets (Huang *et al.* 2016; Foster *et al.* 2012; Fernandez-Garvalho *et al.* 2015; Richards *et al.* 2012). In an Atlantic tuna fishery, the use of circle-hooks did not reduce the by-catch of leatherback sea turtles, but did significantly increase the catch of both swordfish and tuna (Huang *et al.* 2016). In another Atlantic-based study, circle-hooks reduced both leatherback and swordfish catches (Fernandez-Garvalho *et al.* 2015). In the northwest Atlantic, the use of circle-hooks and mackerel bait individually did lessen the by-catch of both leatherback and loggerhead sea turtles, however circle-hooks with squid bait negatively impacted swordfish catch and mackerel bait paired with either a J-hook or a circle-hook reduced tuna catches (Foster *et al.* 2012). Richards *et al.* (2012)found no difference in either catch or bycatch rates between circle-hooks and J-hooks. Moreover, mandatory use of circle-hooks in the U.S. Atlantic pelagic long line fleet has not reduced the rate loggerhead by-catch in this fishery (Fairfield-Walsh & Garrison 2007; Fairfield-Walsh & Garrison 2008; Garrison *et al.* 2009; Garrison & Stokes 2014; Finkheiner *et al.* (2011).

On the other hand, one known advantage of the circle-hook is an increase in the post-hooking survivability of turtles (Finkheiner *et al.* 2011). largely due to the reduction of deeply ingested hooks and mouth/beak hooking events in loggerhead and leatherback sea turtles, respectively (Stokes *et al.* 2012; Pargia 2012:Gilman *et al.* 2007). This shift in the location of hooks on sea turtles lessens the amount of line left trailing after release, which has been identified as the “most dangerous part of the gear” (Pargia 2012). and increases the probability of total gear removal (Stokes *et al.* 2012). It is for this reason that 90% of the Atlantic Canadian swordfish fishing fleet had voluntarily switched to using the circle-hook exclusively by 2010 (DFO 2010). Another method that has been investigated for mitigating turtle by-catch is hooking bait once rather than threading bait, which is the traditional baiting technique. The idea behind this is that sea turtles will spend less time at a baited hook if the bait is more easily removed, thus reducing the likelihood of a hooking incident. Single hooked bait has not been shown to impact sea turtle by-catch and does appear to negatively impact tuna catch rates compared to threading bait (Richards *et al.* 2012). thus is not currently an economically viable option for turtle by-catch mitigation.

## 3. Current Mitigation in the Canadian Pelagic Long Line Fishery

The Nova Scotia Swordfishermen’s Association (NSSA) is the principle organizer of a large bulk of the current turtle by-catch mitigation methods in the Canadian long line fleet (Atlantic Leatherback Turtle Recovery Team 2006; DFO 2010; DFO 2013). A majority of the swordfish-directed long line fleet has been using circle hooks since 1995 to increase the post-release survivability of hooked turtles (Atlantic Leatherback Turtle Recovery Team 2006). With funding from the Habitat Stewardship Program, the NSSA was been able to develop a Code of Conduct for Responsible Sea Turtle Handling and Mitigating Measures in 2003 that includes the avoidance of areas with high sea turtle by-catch, the use of lines long enough to allow captured turtles to reach the surface, and guidelines for the proper handling and safe release of sea turtles (DFO, 2010; DFO 2013). From 2003-2004, the NSSA purchased de-hooking and line-cutting kits for all vessels in the swordfish long line fleet and a fisherman from each vessel had received certification for the use of these kits by 2008 (DFO 2010). A number of mitigation methods outlined in the Code of Conduct have been implemented in the fishery through terms and conditions of licenses, including increased observer coverage, the use of vessel monitoring systems (VMS), the use of safe turtle handling protocols and kits, and the mandatory use of circle hooks starting in 2012 (DFO 2010: DFO 2013).

The NSSA’s initiative to minimize the impact of the long line fishery on the population recovery of Atlantic sea turtles displays the desire of the fishermen to create a sustainable fishery. However, all of the mitigation methods currently in use, save voluntary fleet communication between vessels to identify areas of high turtle by-catch, do not reduce sea turtle incidents, the methods rather lessen the impact of a by-catch event on sea turtles. Since the post-release mortality of sea turtles is unknown, loggerhead and leatherback populations may still be under a great deal of stress from fishing gear in critical foraging habitats. In addition, the negative economic impact of turtle by-catch on fishermen (e.g. loss of hooks, loss of bait, reduced of fishing opportunities, loss of the time spent removing gear and assessing the nature of hooking incidents) has not been alleviated by current mitigation methods. Future mitigation policies should focus on reducing the rate of turtle interactions within the long line fishery.

## 4. Suggestions

There is variability in the effectiveness of by-catch mitigation methods among oceanic regions and among species-specific fisheries within a single region, thus the efficacy and economic viability of various methods should be assessed through experimental fisheries in a region of interest prior to further implementation (FAO 2009). The research and development of fishing practices aimed at reducing bycatch ought to be carried out through collaboration between fisheries managers, scientists, and active fishermen in order to maximize compliance among vessels upon implementation in the commercial fishery (Cox *et al.* 2007). Therefore, the following suggestions should be evaluated through experimentation on active long line vessels in Canadian waters prior to being considered for application via changes in the terms and conditions of fishing licenses.

### 4.1 Temperature-based By-catch "Hotspot" Avoidance

Fishermen should be provided with up-to-date thermal maps of Canadian surface waters so as to assist in the identification of areas that have a high likelihood of sea turtle interactions occurring. The Code of Conduct for Responsible Sea Turtle Handling and Mitigating Measures includes avoiding areas of high turtle abundance as one of its procedures (DFO 2013) and Brazner & McMillan (2008) recommend that setting gear on the cold side of an oceanic front will separate fishing effort and foraging sea turtles in space without negatively impacting target catch. The TurtleWatch program in the Hawaiian-based long line fishery has confirmed that sea surface temperature can be a reliable tool for assessing the likelihood of sea turtle interactions in a region (Howell *et al.* 2008). therefore a similar program should be developed and made readily available for the Canadian Pelagic Long Line Fleet to aid fishermen in the classification of areas with a high risk for turtle bycatch. Fishermen will be able to review the thermal properties of potential fishing grounds on their steam out to sea and infer the ideal range at which to deploy their gear. It is in the best interest of the fishermen to reduce sea turtle by-catch through voluntary avoidance so as to maximize fishing selectivity in the short term and avoid the implementation of more restrictive measures (e.g. by-catch caps, reduced fishing effort, shorter seasons) in the future.

### 4.2 Restrict Tuna-directed Long Line Sets to Depths Greater than 40 meters

In the Canadian Pelagic Long Line Fleet, sets directed at tuna historically have a higher turtle by-catch rate than those targeting swordfish (Paul *et al.* 2010; Brazner & McMillan 2008). Thus, establishing mitigation methods specifically for the long line tuna fleet may be an efficient strategy to reduce the overall impact of Canadian fisheries on sea turtle populations. Limiting the depth of Canadian tuna fishing operations to waters deeper than 40 meters would separate sea turtles and baited hooks vertically in space (Bailey *et al.* 2012; Wallace *et al.* 2015; Bolton 2003) and reduce sea turtle interactions, given that by-catch by Canadian long lines takes place almost exclusively in the top 40 meters of the water column (Brazner & McMillan 2008). Tuna are traditionally fished during the day, which happens to overlap with the deepest extent of the fish’s diel vertical migration (Lam *et al.* 2014; Dagorn *et al.* 2000). Adult tuna in the northwest Atlantic frequent an average depth of approximately 200 meters during the day and move up the water column to about 45 meters at night (Lam *et al.* 2014). shadowing the vertical migration of their prey species (Vaske *et al.* 2012). Tuna primarily forage at sub-thermocline depths during the day (Vaske *et al.* 2012). Therefore, this vertical shift in fishing effort is expected to have negligible impacts on tuna catch.

Furthermore, this mitigation technique could assist fishermen in selectively fishing for larger adult tuna while minimizing the catch of juveniles. Juvenile tuna generally remain in the top 40 meters of the water column (Brill *et al.* 2002). displaying different diel vertical patterns than those of larger tuna (Dagorn *et al.* 2000). The size of a tuna determines the rate at which muscle temperatures change with variations in surround water temperatures, with larger tuna cooling more slowly, permitting them to dive to and remain in deeper waters (Dagorn *et al.* 2000). It follows that fishing at depths greater than 40 meters could increase the catch weight per unit of effort of tuna species targeted by the Canadian Pelagic Long Line Fleet. On the other hand, swordfish fishing vessels set their gear at night in shallow waters (FAO 2009) to exploit the distinct diel vertical migration of swordfish that forage in surface waters during the nighttime (Dewar *et al.* 2011). Thus, a shift in fishing operations to deeper depths would not be an economically viable mitigation technique for the long line fleet directed towards swordfish.

## 5 Conclusion

Atlantic Canadian waters pose a high threat to the population recovery of loggerhead and leatherback sea turtles due to the presence of fishing operations in critical northern habitat. The pelagic long line fleet has been identified as the most significant source of sea turtle mortality in Canadian waters. Although mortality on the line is extremely rare, the post-release mortality of sea turtles captured in long line gear is expected to be high due to the adverse affects of gear that remains attached, primarily fishing line. Canadian fishermen have initiated the current sea turtle by-catch mitigation through voluntary measures to increase the likelihood of survival for sea turtles released from long line gear. Since the post-release mortality of sea turtles is not known, the effectiveness of these measures cannot currently be evaluated. Future mitigating measures should focus on reducing the rate of sea turtle by-catch in the long line fleet in order to minimize the impact of the fishery on sea turtles and lessen the economic burden of by-catch events on the fishermen. Two methods that would likely be effective in Canadian waters include providing fishermen with up-to-date thermal maps of the sea surface to avoid setting gear in areas with a high likelihood of sea turtle presence and lowering the depth of fishing operations that target tuna. These measures should be tested in experimental fisheries prior to implementation to assure effectiveness and economic viability.

## Acknowledgements

Thanks to Jim Williams for helping organize this work and providing guidance throughout the research process. Carolyn Finkle and Averell Sherker reviewed drafts of this paper, which impacted the final version greatly.

## References

Arendt, M.D., Boynton, J., Schwenter, J.A., Byrd, J.I., Segars, A.L., Whitaker, J.D., Parker, L., Owens, D.W., Blanvillain, G.M., Quattro, J.M., and Roberts, M.A. 2012. Spatial clustering of loggerhead sea turtles in coastal waters of the NW Atlantic Ocean: implications for management surveys. Endang. Species Res. 18: 219–231

Atlantic Leatherback Turtle Recovery Team 2006. Recovery strategy for leatherback turtle (*Dermochelys coriacea*) in Atlantic Canada. SARA Ser. Fisheries and Oceans Canada, Ottawa

Bailey, H., Fossette, S., Bograd, S.J., Shillinger, G.L., Swithenbank, A.M., Georges, J-Y., Gaspar, P., Strómberg, K.H.P., Paladino, F.V., Spotila, J.R., Block, B.A., and Hays, G.C. 2012. Movement patterns for a critically endangered species, the leatherback turtle (*Dermochelys coriacea*), linked to foraging success and population status. PLoS ONE 7(5)

Baum, J.K., Myers, R.A., Kehler, D.G., Worm, B., Harley, S.J., and Doherty, P.A. 2003. Collapse and conservation of shark populations in the northwest Atlantic. Sci. 299: 389–392

Beverly, S., Curran, D., Musyl, M., and Molony, B. 2009. Effects of eliminating shallow hooks from tuna longline sets on target and non-target species in the Hawaii-based pelagic tuna fishery. Fish. Res. 96: 281–288

Bolton, A.B. 2003. Active swimmers-passive drifters: the oceanic juvenile stage of loggerheads in the Atlantic system. In: Loggerhead sea turtle. Eds: Bolton, A.B. and Witherington, B.E.. Smithson. Instit. Press, Washington, D.C., p. 63–78

Bowan, B.W., Bass, A.L., Soares, L., and Toonen, R.J. 2005. Conservation implications of complex population structure: lessons from the loggerhead turtle (*Caretta caretta*). Mol. Ecol. 14: 2389–2402

Bowan, B.W. and Karl, S.A. 2007. Population genetics and phylogeography of sea turtles. Mol. Ecol. 16: 4886–4907

Braun-McNeill, J., Sasso, C.R., Epperly, S.P., and Rivero, C. 2008. Feasibility of using sea surface temperature imagery to mitigate cheloniid sea turtle-fishery interactions off the coast of northeastern USA. Endang. Species. Res. 5: 257–266

Brazner, J.C. and McMillan, J. 2008. Loggerhead turtle (*Caretta caretta*) by-catch in Canadian pelagic long line fisheries: relative importance in the western North Atlantic and opportunities for mitigation. Fish. Res. 91(2–3): 310–324

Brill, R., Lutcavage, M., Metzger, G., Bushnell, P., Arendt, M., Lucy, J., Watson, C., and Foley, D. 2012. Horizontal and vertical movements of juvenile Bluefin tuna (*Thunnus thynnus*) in relation to oceanographic conditions of the western North Atlantic, determined with ultrasound telemetry. Fish. Bull. 100:155–167

Casale, P. and Tucker, A.D. 2015. Caretta caretta. The IUCN red list of threatened species

Casey, J.P., James, M.C., Williard, A.S. 2014. Behavioral and metabolic contributions to thermoregulation in freely swimming leatherback turtles at high latitudes. J.Exp. Biol. 217: 2331–2337

Chaloupka, M., Parker, D., and Balazs, G. 2004. Modeling post-release mortality of loggerhead sea turtles exposed to the Hawaiian-based pelagic long line fishery. Mar. Ecol. Prog. Ser. 280: 285–293

COSEWIC 2010. COSEWIC assessment and status report on the loggerhead sea turtle *Caretta caretta* in Canada. Committee on the Status of Endangered Wildlife in Canada, Ottawa

COSEWIC 2012. COSEWIC assessment and status report on the leatherback sea turtle *Dermochelys coriacea* in Canada. Committee on the Status of Endangered Wildlife in Canada. Ottawa

Cox, T.M., Lewison, R.L., Zydelis, R., Crowder, L.B., Safina, C., and Read, A.J. 2007. Comparing effectiveness of experimental and implemented by-catch reduction measures: the ideal and the real. Cons. Biol. 21(5): 1155–1164

Dagorn, L., Bach, P., and Josse, E. 2000. Movement patterns of large bigeye tuna (*Thunnus obesus*) in the open ocean, determined using ultrasonic telemetry. Mar. Biol. 136: 361–371

Davoren, G.K. 2013. Distribution of marine predator hotspots explained by persistent areas of prey. Mar. Bio. 160(12): 3043–3058

Dewar, H., Prince, E.D., Musyl, M.K., Brill, R.W., Sepulveda, C., Luo, J., Foley, D., Orbensen, E.S., Domeier, M.L., Nasby-Lucas, N., Snodgrass, D., Laurs, R.M., Hoolihan, J.P., Block, B.A., and McNaughton, L.M. 2011. Movements and behaviors of swordfish in the Atlantic and Pacific Oceans examined using pop-up satellite archival tags. Fish. Oceanog. 20: 219–241

DFO 2010. Atlantic loggerhead conservation action plan. DFO Can. Sci. Advis. Sec. Proceed. Ser. 2010

DFO 2013. Canadian Atlantic swordfish and other tunas. DFO Can. Sci. Advis. Sec. Proceed. Ser. 2013

Dodge, K.L., Galuardi, B., Miller, T.J., and Lutcavage, M.E. 2014. Leatherback turtle movements, dive behavior, and habitat characteristics in ecoregions of the northwest Atlantic Ocean. PLoS ONE 9(3)

Dunn, D.C., Boustany, A.M., and Halpin, P.N. 2011. Spatio-temporal management of fisheries to reduce by-catch and increase fishing selectivity. Fish and Fisheries 12: 110–119

Edgar, G.J., Russ, G.R., and Babcock, R.C 2007. Marine Protected Area In: Marine ecolog Ed: Sean Connell. Oxford University Press, South Melbourne

Ehrhart, L., Redfoot, W., Bagley, D., and Mansfield, K. 2014. Long-term trends in loggerhead (*Caretta caretta*) nesting and reproductive success at an important western Atlantic rookery. *Chelo. Cons. Biol.* 13: 173–181

Epperly, S.P. 2003. Fisheries-related mortality and turtle excluder devices (TEDs) In: The biology of sea turtles, volume I. Eds: Lutz, P.J., Musick, J.A., and Wyneken, J.. CRC Press, Boca Raton, Florida, p. 339–353

Fairfield-Walsh, C. and Garrison, L.P. 2007. Estimated by-catch of marine mammals and turtles in the U.S. Atlantic pelagic long line fleet during 2006. NOAA Tech. Memor. NMFS-SEFSC 560: 1–58

Fairfield-Walsh, C. and Garrison, L.P. 2008. Estimated by-catch of marine mammals and turtles in the U.S. Atlantic pelagic long line fleet during 2007. NOAA Tech. Memor. NMFS-SEFSC 572:1–63

FAO 2009. Guidelines to reduce sea turtle mortality in fishing operations. FAO, Rome

Fernández-Álamo, M.A. and Färber-Lorda, J. 2006. Zooplankton and the oceanography of the eastern tropical Pacific: a review. Prog. Oceano. 69: 318–359

Fernandez-Carvalho, J., Coelho, R., Santos, M.N., and Amorim, S. 2015. Effects of hook and bait in a tropical northeast Atlantic pelagic long line fishery: part II-target, by-catch, and discard fish. Fish. Res. 164: 312–321

Finkbeiner, E.M., Wallace, B.P., Moore, J.E., Lewison, R.L., Crowder, L.B., and Read, A.J. 2011. Cumulative estimates of sea turtle by-catch and mortality in USA fisheries between 1990 and 2007. Biol. Cons. 144: 2719–2727

Fossette, D., Girard, C., López-Mendilaharsu, M., Miller, P., Domingo, A., Evans, D., Kelle, L., Plot, V., Prosdocimi, L., Verhage, S., Gaspar, P., and Georges, J-Y. 2010. Atlantic leatherback migratory paths and temporary residence areas. PLoS ONE 5(11)

Foster, D.G., Epperly, S.P., Shah, A.K., and Watson, J.W. 2012. Evaluation of hook and bait type on the catch rates in the western North Atlantic Ocean pelagic long line fishery. Bull. Mar. Sci. 88(3): 529–545

Gardner, B., Sullivan, P.J., Morreale, S.J., and Epperly S.P. 2008. Spatial and temporal statistical analysis of bycatch data: patterns of sea turtle by-catch in the North Atlantic. Can.J. Fish. Aquat. Sci. 65: 2461–2470

Garrison, L.P., Stokes, L., and Fairfield, C. 2009. Estimated by-catch of marine mammals and sea turtles in the U.S. Atlantic pelagic long line fleet during 2008. NOAA Tech. Memor. NMFS-SEFSC 591

Garrison, L.P. and Stokes, L. 2014. Estimated by-catch of marine mammals and sea turtles in the U.S. Atlantic pelagic long line fleet during 2013. NOAA Tech. Memor. NMFS-SEFSC 667:1–61

Game, E.T., Grantham, H.S., Hobday, A.J., Pressey, R.L., Lombard, A.T., Beckley, L.E., Gjerde, K., Bustamante, R., Possingham, H.P., and Richardson, A.J. 2009. Pelagic protected areas: the missing dimension in ocean conservation. TREE 24(7): 360–369

Gilman, E., Zollett, E., Beverly, S., Nakano, H., Davis, K., Shiode, D., Dalzell, P., Kinan-kelly, I. 2006. Reducing sea turtle by-catch in pelagic long line fisheries. Fish and Fisheries 7(1): 2–23

Gilman, E., Kobayashi, D., Swenarton, T., Brothers, N., Dalzell, P., and Kinan-Kelly, I. 2007. Reducing sea turtle interactions in the Hawaiian-based long line swordfish fishery. Biol. Cons. 139(1–>2): 19–28

Greer, A.E. 1973. Anatomical evidence for a counter-current heat exchanger in the leatherback turtle (*Dermochelys coriacea*). Nature 244(5412): 181

Hamelin, K.M., Kelley, D.E., Taggart, C.T., and James, M.C. 2014. Water mass characteristics and solar illumination influence leatherback turtle dive patterns at high latitudes. Ecol. Soc. Amer. 5(2): 1–20

Harris, L.E., Gross, W.E., Smedbol, R.K., and Bondt, L.H. 2010. Loggerhead turtles (*Caretta caretta*) in Atlantic Canada: biology, status, recovery potential, and measures for mitigation. Can. Sci. Adv. Sec. Res. Doc. 89

Hays, G.C., Broderick, A.C., Godley, B.J., Luschi, P., and Nichols, W.J. 2003. Satellite telemetry suggests high levels of fishing-induced mortality in marine turtles. Mar. Ecol. Prog. Ser. 262: 305–309

Hays, G.C. 2008. Sea turtles: a review of some key recent discoveries and remaining questions. J.Exp. Mar. Biol. Ecol. 356:1–7

Hochscheid, S., Bentivegna, F., Bradai, M.N., and Hays, G.C 2007. Overwintering behavior in sea turtles: dormancy is optional. Mar. Ecol. Prog. Ser. 340: 287–298

Houghton, J.D.R., Doyle, T.K., Wilson, M.W., Davenport, J., and Hays, G.C. 2006. Jellyfish aggregations and leatherback turtle foraging patterns in a temperate coastal environment. Ecol. 87(8): 1967–1972

Howell, E.A., Kobayashi, D.R., Parker, D.M., Balazs, G.H., and Polovina, J.J. 2008. TurtleWatch: a tool to aid in the bycatch reduction of loggerhead turtle *Caretta caretta* in the Hawaii-based pelagic longline fishery. Endang. Species Res. 5: 267–278

Howell, E.A., Hoover, A., Benson, S.R., Bailey, H., Polovina, J.J., Seminoff, J.A., and Dutton, P.H. 2015. Enhancing the TurtleWatch product for leatherback sea turtles, a dynamic habitat model for ecosystem-based management. Fish. Oceanogr. p. 1–12

Huang, H.W., Swimmer, Y., Bigelow, K., Gutierrez, A., and Foster, D.G. 2016. Influence of hook type on catch of commercial and by-catch species in an Atlantic tuna fishery. Mar. Poll 65: 68–75

IAC Sectretariat 2004. Inter-American convention for the protection and conservation of sea turtles-an introduction, September 2004

James, M.C., and Herman, T.B. 2001. Feeding of *Dermochelys coriacea* on medusea in the northwest Atlantic. Chelo. Cons. Bio. 4: 202–205

James, M.C. and Mrosovsky, N. 2004. Body temperatures of leatherback turtles (*Dermochelys coriacea*) in temperate waters off of Nova Scotia, Canada. Can.J. Zool. 82: 1302–1306

James, M.C., Ottensmeyer, C.A., and Meyers, R.A. 2005a. Identification of high-use habitat and threats to leatherback sea turtles in northern waters: new directions for conservation. Ecol. Lett. 8: 195–201

James, M.C., Meyers, R.A., and Ottensmeyer, C.A. 2005b. Behaviour of leatherback sea turtles *Dermochelys coriacea*, during the migratory cycle. Proc. R. Soc. B 272:1547–1555

James, M.C., Sherrill-Mix, S.A., and Myers, R.A. 2007. Population characteristics and seasonal migrations of leatherback sea turtles at high latitudes. Mar. Ecol. Prog. Ser. 337: 245–254

James, M.C., Ottensmeyer, C.A., Eckert, S.A., and Myers, R.A. 2006. Changes in diel diving patterns accompany shifts between northern foraging and southward migration in leatherback turtles. Can.J. Zool. 84: 754–765

Jones, T.T., Bostrom, B.L., Hastings, M.D., Van Houtan, K.S., Pauly, D., and Jones, D.R. 2012. Resource requirements of the Pacific leatherback turtle population. PLoS ONE 7(10)

Lam, C.H., Galuardi, B., and Lutcavage, M.E. 2014. Movements and oceanographic associations of bigeye tuna (*Thunnus obesus*) in the northwest Atlantic. Can.J. Fish. Aquat. Sci. 71: 1–15

Lewison, R.L., Freeman, S.A., and Crowder, L.B. 2004. Quantifying the effects of the fisheries on the threatened species: the impacts of pelagic long lines on the loggerhead and leatherback sea turtle. Ecol. Lett. 7(3): 221–231

Lewison, R.L., Soykan, C.U., and Franklin, J. 2009. Mapping the bycatch seascape: multispecies and multi-scale spatial patterns of fisheries bycatch. Ecol. Appl. 19:920–930

Lewison, R., Wallace, B., Alfaro-Shigueto, J., Mangel, J.C., Maxwell, S.M., and Hazen, E.L. 2013. Marine turtles: lessons learned from decades of research and conservation. In: The biology of sea turtles, volume II. Eds: Wyneken, J., Lohmann, K.J., and Musick, J.A. CRC Press, Boca Raton, London, New York, p. 329–351

Liberal Party of Canada 2016. Real change: protecting our oceans. https://www.liberal.ca/realchange/trudeau-announces-plan-to-protect-canadas-oceans/

Mansfield, K.L., Saba, V.S., Keinath, J.A., and Musick, J.A. 2009. Satellite tracking reveals a dichotomy in migration strategies among juvenile loggerhead turtles in the northwest Atlantic. Mar. Biol. 156: 2555–2570

McAlpine, D.F., James, M.C., Lein, J., and Orchard, S.A. 2007. Status and conservation of marine turtles in Canadian waters. In: Ecology, conservation, and status of reptiles in Canad. Eds: Seburn, C.N.L. and Bishop, C.A.. Herp. Cons. 2: 85–112

McMahon, C.R. and Hays, G.C. 2006. Thermal niche, large-scale movements and implications of climate change for a critically endangered marine vertebrate. Glob. Chan. Biol. 12: 1330–1338

Murawski, S.A., Wigley, S.E., Fogarty, M.J., Rago, P.J., and Mountain, D.G. 2005. Effort distribution and catch patterns adjacent to temperate MPAs. J.Mar. Sci. 62(6): 1150–1167

Murray, K.T. 2009. Characteristics and magnitude of sea turtle by-catch in U.S. midAtlantic gillnet gear. Endang. Species Res. 8: 211–224

Narazaki, T., Sato, K., and Miyazaki, N. 2015. Summer migration to temperate foraging habitats and active winter diving of juvenile loggerhead turtles *Caretta caretta* in the western North Pacific. Mar. Biol. 162(6): 1251–1263

NOAA Fisheries 2014. Loggerhead sea turtle (*Caretta caretta*)

O’Keefe, C.E., Cadrin, S.X., and Stokesbury, K.D.E. 2013. Evaluating effectiveness of time/area closures, quota/caps, and fleet communication to reduce fisheries bycatch. ICES J, of Mar. Sci. 71(5): 1286–1297

Pargia, M.L. 2012. Hooks and sea turtles: a veterinarian’s perspective. Bull. Mar. Sci. 88(3): 731–741

Parker, D.M., Cooke, W.J., and Balazs, G.H. 2005. Diet of oceanic loggerhead sea turtles (*Caretta caretta*) in the central North Pacific. Fish. Bull. 103: 142–152

Paul, S.D., Hanke, A., Smith, S.C., and Neilson, J.D. 2010. An examination of loggerhead sea turtle (*Caretta caretta*) encounters in the Canadian swordfish and tuna long line fishery, 2002–2008. Can. Sci. Adv. Sec. Res. Doc.

Polovina, J.J., Howell, E., Parker, D.M., and Balazs, G.H. 2003. Dive-depth distribution of loggerhead (*Caretta caretta*) and olive ridley (*Lepidochelys olivacea*) sea turtles in the central North Pacific: might deep long line sets catch fewer turtles?. Fish. Bull. 101(1): 189–193

Putman, N.F. and Mansfield, K.L. 2015. Direct evidence of swimming demonstrates active dispersal in the sea turtle “lost years”. Curr. Biol. 25:1–7

Richards, P.M., Epperly, S.P., Watson, J.W., Foster, D.G., Bergmann, C.E., and Beideman, N.R. 2012. Can circle hook offset combined with baiting technique affect catch and by-catch in pelagic long line fisheries. Bull. Mar. Sci. 88(3): 589–603

Roe, J.H., Morreale, S.J., Paladino, F.V., Shillinger, G.L., Benson, S.R., Eckert, S.A., Bailey, H., Tomillo, P.S., Bograd, S.J., Eguchi, T., Dutton, P.H., Seminoff, J.A., Block, B.A., and Spotila, J.R. 2014. Predicting by-catch hotspots for endangered leatherback turtles on longlines in the Pacific Ocean. Proc. R. Soc. B 281(1777)

Ryder, C.E., Conant, T.A., and Schroeder, B.A. 2006. Report of the workshop on marine turtle long line post-interaction mortality. US Dep. Commerce, NOAA.Tech. Memor. NMFS-F/OPR 29: 1–36

Sales, G., Giffoni, B.B., Fielder, F.N., Azevedo, V.G., Kotas, J.E., Swimmer, Y., and Bugoni, L. 2010. Circle hook effectiveness for the mitigation of sea turtle bycatch and capture of target species in a Brazilian pelagic long line fishery. Aqua. Cons.: Mar. Freshwat. Ecosyst. 20: 428–436

Santos, M.N., Coelho, R., Fernandez-Carvalho, J., and Amorim, S. 2013. Effects of 17/0 circle hooks and bait on sea turtles by-catch in a southern Atlantic swordfish longline fishery. Aqua. Cons.: Mar. Freshwat. Ecosyst. 23(5): 732–744

Sasso, C.R. and Epperly, S.P. 2007. Survival of pelagic juvenile loggerhead turtles in the open ocean. J. Wildl. Manage. 71(6): 1830–1835

Sherrill-Mix, S.A., James, M.C., and Myers, R.A. 2008. Migration cues and timing in leatherback sea turtles. Behav. Ecol. 19(2): 231–236

Shwartz, F.J. 1978. Behavioral and tolerance responses to cold water temperatures by three species of sea turtles (Reptilia, Cheloniidae) in North Carolina. Florida Mar. Res. Publ. 33: 16–18

Smolowitz, R.J., Patel, S.H., Haas, H.L., and Miller, S.A. 2015. Using a remotely operated vehicle (ROV) to observe loggerhead sea turtle (*Caretta caretta*) behavior on foraging grounds off the mid-Atlantic United States. J. Exp. Mar. Biol. Ecol. 471: 84–91

Stokes, L.W., Epperly, S.P. and McCarthy, K.J. 2012. Relationship between hook type and hooking location in sea turtles incidentally captured in the United States Atlantic pelagic long line fishery. Bull. Mar. Sci. 88: 703–718

Swimmer, Y., Arauz, R., McCracken, M., McNaughton, L., Ballestero, J., Musyl, M., Bigelow, K., and Brill, R. 2006. Diving behavior and delayed mortality of olive ridley sea turtles *Lepidochelys olivacea* after their release from long line fishing gear. Mar. Ecol. Prog. Ser. 323: 253–261

Swimmer, Y., Empey Campora, C., McNaughton, L., Musyl, M., and Pargia, M. 2014. Post-release mortality estimates of loggerhead sea turtles (*Caretta caretta*) caught in the pelagic long line fisheries based on satellite data and hooking location. Aqua. Cons.: Mar. Freshwat. Ecosyst.

Vaske, T., Travassos, P.E., Hazin, F.H.V., Tolotti, M.T., and Barbosa, T.M. 2012. Forage fauna in the diet of bigeye tuna (*Thunnus obesus*) in the western tropical Atlantic Ocean. Braz.J. Oceanogr. 60(1): 89–97

Wallace, B.P., DiMatteo, A.D., Hurley, B.J., Finkbeiner, E.M., Bolton, A.B., Chaloupka, M.Y., Hutchinson, B.J., Abreu-Grobois, F.A., Amorocho, D., Bjorndal, K.A., Bourjea, J., Bowen, B.W., Briseño-Dueñas, R., Casale, P., Choudhury, B.C., Costa, A., Dutton, P.H., Fallabrino, A., Girard, A., Girondot, M., Godfrey, M.H., Hamann, M., López-Mendilaharsu, M., Marcovaldi, M.A., Mortimer, J.A., Musick, J.A., Nel, R., Pilcher, N.J., Seminoff, J.A., Troëng, S., Witherington, B., and Mast, R.B. 2010. Regional Management Units for marine turtles: a novel framework for prioritizing conservation and research across multiple scales. PLoS ONE 5(12)

Wallace, B.P., DiMatteo, A.D., Bolton, A.B., Chaloupka, M.Y., Hutchinson, B.J., Abreu-Grobois, F.A., Mortimer, J.A., Seminoff, J.A., Amorocho, D., Bjorndal, K.A., Bourjea, J., Bowen, B.W., Briseño-Dueñas, R., Casale, P., Choudhury, B.C., Costa, A., Dutton, P.H., Fallabrino, A., Finkbeiner, E.M., Girard, A., Girondot, M., Hamann, M., Hurley, B.J., López-Mendilaharsu, M., Marcovaldi, M.A., Musick, J.A., Nel, R., Pilcher, N.J., Troëng, S., Witherington, B., and Mast, R.B. 2011. Global conservation priorities for marine turtles. PLoS ONE 6(9)

Wallace, B.P., Kot, C.Y., DiMatteo, A.D., Lee, T., Crowder, L.B., and Lewison, R.L. 2013. Impacts of fisheries by-catch on marine turtle populations worldwide: toward conservation and research priorities. Ecol. Soc. Amer. 4(3): 1–49

Wallace, B.P., Zolkewitz, M., and James, M.C. 2015. Fine-scale foraging ecology of leatherback turtles. Front. Ecol. Evol. 3:15

Watson, J.W., Epperly, S.P., Shah, A.K., and Foster, D.G. 2005. Fishing Methods to reduce sea turtle mortality associated with pelagic longlines. Can. J. Fish. Aquat. Sci. 62: 965–981

Witherington, B., Kubilis, P., Brost, B., and Mevlan, A. 2009. Decreasing annual nest counts in a globally important loggerhead sea turtle population. Ecol. Appl. 19(1): 30–54

Witt, M.J., Broderick, A.C., Johns, D.J., Martin, C., Pennose, R., Hoogmoed, M.S., and Godley, B.J. 2007. Prey landscapes help identify potential foraging habitats for leatherback turtles in the northeast Atlantic. Mar. Ecol. Prog. Ser. 337: 231–243

Witt, M.J., Hawkes, L.A., Godfrey, M.H., Godley, B.J., and Broderick, A.C. 2010. Predicting the impacts of climate change on a globally distributed species: the case of the loggerhead turtle. J. Exp. Biol. 213: 901–911

Zug, G.R., Vitt, L.J., and Caldwell, J.P. 2001. Herpetology: introductory biology of amphibians and reptiles, second edition. Academ. Press, San Diego, p. 1–630

